# Global context rapidly shapes sensory responses in V1

**DOI:** 10.64898/2026.01.07.698143

**Authors:** Darcy S. Peterka, Fumiyasu Imai, Jordan M. Ross, Georgia Bastos, Molly Hornick, Lital Rachmany, Connor G. Gallimore, Adam Hockley, Jordan P. Hamm

## Abstract

Context modulates sensory processing in the cerebral cortex by suppressing responses to expected stimuli and enhancing responses to unexpected ones. Recent proposals argue that early sensory areas such as primary visual cortex (V1) are shaped only by local context, including recent stimulus history, whereas modulation by global context, such as learned temporal structure, is present exclusively in higher cortical areas. This view is incompatible with predictive coding theories. To directly dissociate local and global contextual influences, we used a global/local oddball paradigm in which mice viewed five-item sequences. Across conditions, sequence structure was held constant while stimulus identity and predictability were selectively manipulated, allowing the isolation of response modulations due to local deviance, global expectation, and stimulus repetition independently. In the canonical sequence (AAAA–B), B is locally deviant but globally predictable. Using two-photon calcium imaging and LFP recordings in mouse V1, we found that global predictability abolished context modulation: responses to B were equivalent to those evoked by a random sequence control (e.g., CDEAB). This effect emerged rapidly, after only <10 sequence repetitions, demonstrating fast learning of global structure. When the stimulus was globally deviant, either by replacing B with a novel stimulus (AAAA–C) or by presenting B unpredictably in a standard oddball paradigm, V1 exhibited robust response enhancement. These effects required feedback from anterior cingulate area (ACa), establishing a causal role for higher cortical circuits in conveying global predictions to V1. Strikingly, when an additional A replaced B (AAAA–A), responses were strongly suppressed despite global deviance, indicating that stimulus-specific adaptation may constrain the expression of global prediction error signals in early sensory cortex.

**Highlights:** Global context rapidly modulates neural responses in V1

Global deviations, but not predictable local deviants, elicit enhanced responses

Downstream brain region ACa is necessary for global context modulation in V1

Standard oddball paradigms involve higher-order contextual modulation

## Introduction

Sensory processing regions of the cerebral cortex integrate incoming input from the periphery within broader context, including concurrent and recent stimuli, cross-modal inputs, learned spatiotemporal regularities, and behavioral state. Predictive processing theories posit that this integration involves suppression of cortical responses to stimuli that are predictable in that context, and amplification of responses to stimuli that deviate from contextual regularities^1–3^. Evidence for such modulation is present as early as primary visual cortex (V1)^4,5^ or primary auditory cortex (A1)^6,7^, but much of this evidence comes from experimental studies using a standard “oddball” sequence, involving a repeated redundant stimulus “A” with a rare deviant stimulus “B” interspersed at random times (e.g. AAAAABAAB).

There is ongoing debate as to whether modulation of V1 responses can reflect higher-level expectations, stemming from global probabilities encoded over longer time scales, or simply the consequences of local adaptation to immediately preceding stimuli - i.e., purely local context^8^. The latter would represent a challenge to classical predictive processing theory – at least insofar as it holds for V1. A recent report concluded that enhanced V1 responses observed during oddball paradigms reflect responses to deviation (i.e. “deviance detection”) *only* from local context^9^. Specifically, Westerberg et al used a “global/local” oddball paradigm and found that repeatedly presenting predictable four stimulus sequences such as ”AAAB” to a mouse left large V1 responses to “B” intact (local deviant, global redundant), despite the fact that the “B” stimulus was predictable in the global (i.e., longer term) context. Likewise, no such large responses were in V1 when stimulus “A” replaced the predictable B at the end of the sequence (i.e., “AAAA” local redundant, global deviant), although higher cortical areas appeared to detect this deviation. Authors concluded that a) this conflicts with the notion that global context can modulate processing in V1 and b) refutes standard predictive coding theories, which posit top-down suppression of sensory cortical responses to contextually predicted stimuli. Instead, they propose that oddball responses in V1 arise from stimulus adaptation in feed-forward circuitry. Similarly, in an auditory global/local oddball study, no A1 responses to local redundant, global deviant “AAAAA” stimuli were present across the neural population, though small increases occur in parvalbumin positive interneurons^10^. Here authors suggested that global context may be encoded at higher levels of the cortical hierarchy, with weak top-down signals to sensory cortices.

These conclusions are at odds, however, with two lines of work. One, using local field potential recordings (LFP) in V1 have evinced clear deviance detection when complex sequences designed to circumvent local adaptation (e.g. ABCD) are violated (e.g. ABDC)^11^, yet whether these LFP signals reflect V1 neuronal outputs or distal feedback signals from other brain regions cannot be discerned from LFP alone. Second, other reports studying both V1^4,5^ and A1^7^ neurons directly show that, even during standard oddball paradigms (i.e, B stimuli occurring at random intervals, e.g. AAAAABAAB)., feedback modulation from higher brain regions (e.g. medial prefrontal areas in the mouse) is needed for the full range of context effects – specifically, deviance detection. Importantly, in these standard oddball paradigms^4,5,7^, stimuli B are, indeed, locally deviant just as they are in the AAAB sequences from traditional global/local paradigms, but the timing of stimulus B is not predictable. Thus, in a standard oddball paradigm, B stimuli are both local and global oddballs, complicating comparison between these two types of studies.

One idiosyncrasy of traditional global/local paradigms is that the global deviant in AAA-A is also the locally redundant stimulus^9,10^. That is, replacing the predictable “B” in the AAAB sequence with an “A” creates a condition where neural adaptation (a suppressive effect) and deviance detection (an amplifying effect) are co-occurring yet inverse effects. This complicates interpretations, as our past work has shown that stimulus specific adaptation (response reductions) to the repeated “A” in a standard oddball sequence is rapid and very strong in V1. Also, such stimulus specific adaptation a) is mechanistically distinct from deviance detection, as it depends neither on local SST+ or VIP+ interneurons nor top-down feedback modulation^4,12^, and b) is present earlier and at lower levels of the visual processing hierarchy than deviance detection (i.e. in layer 4 responses and extracellular currents^13^). Likewise, in the auditory system stimulus specific adaptation is present as early as midbrain and thalamus^14^, several synapses upstream from cortex. Thus, it is possible that the failure to identify global deviance detection using traditional global/local paradigms – and thus, also prediction error-like responses – in V1 to the “A” stimulus may stem from the particular stimulus design: that this global deviant was presented to a system that had already strongly adapted to the relevant features (i.e. to the “A”), perhaps even upstream of V1.

Here we sought to replicate and then extend the results reported in Westerberg et al and the theory presented in Gabhart et al ^8^ using a similar “global/local” oddball paradigm, through stimuli designed with two additional changes. 1) We presented a condition where the final stimulus in the sequence is both a global *and* a local deviant (“C”). And 2) we included a random orientation control run, where bursts of 5 stimuli are presented of random orientations (Figure 1B). This second change allowed for comparison of responses to stimulus “A” (90 or 0 degrees), “B” (0 or 90 degrees) and “C” (45 degrees) when they appeared at the end of an unpredictable stimulus chain (e.g. CDAEB; 8 possible orientations, spaced 22.5 degrees) vs when they came at the end of a predictable stimulus chain (AAAA-x).

**Figure 1:**
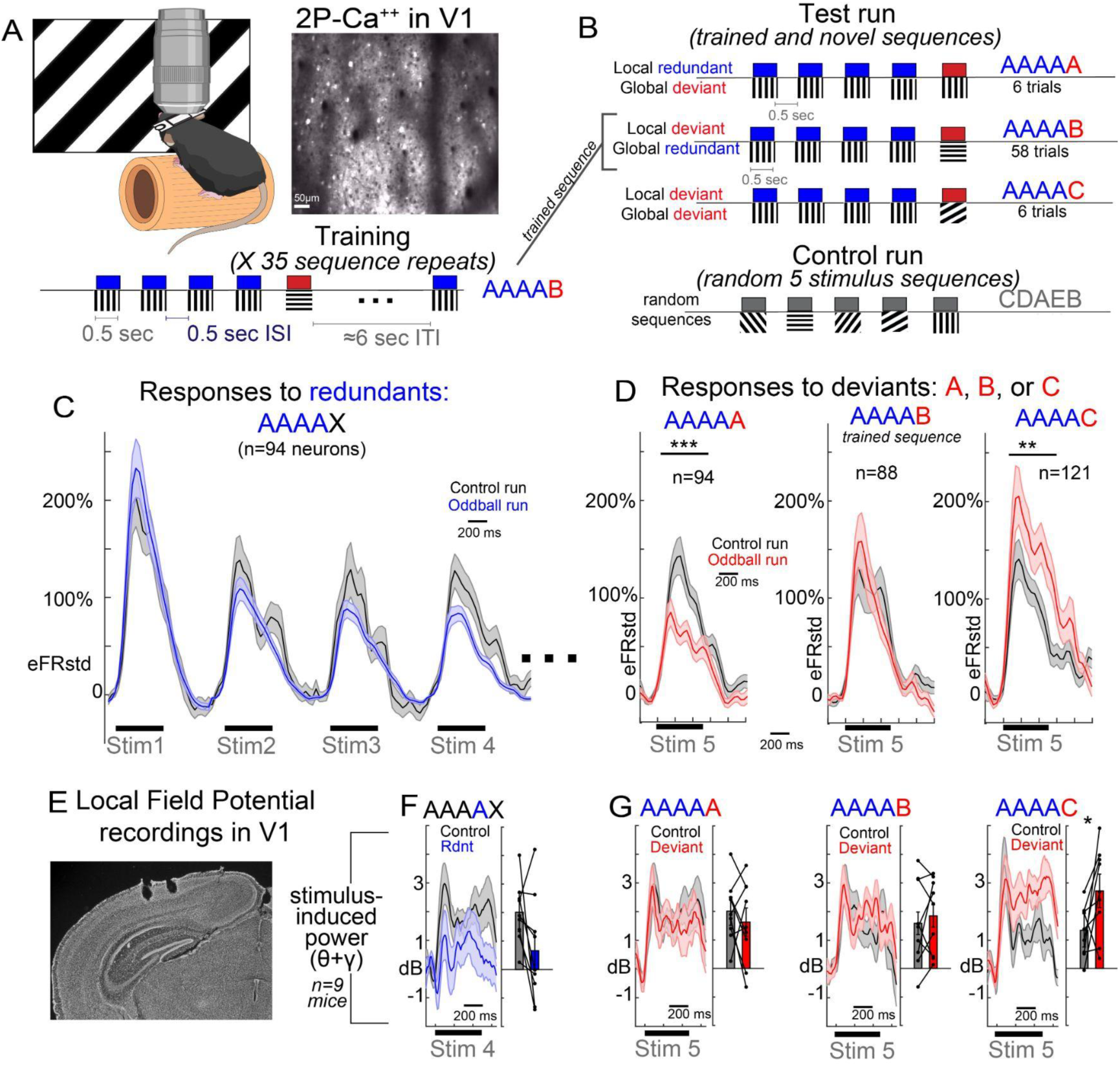
Deviation from global context is detected in V1 but obscured by stimulus specific adaptation. A) Head-fixed mice viewed full field moving oriented grating stimuli while V1 neurons were recorded with two-photon calcium imaging (GCaMP6s). Mice viewed 5 stimulus sequences (4.5 second duration) of AAAAB, presented 35 times for training/habituation. B) Then, after a short (<3 minute) break, mice viewed a test run during which the final “B” in the sequence was occasionally replaced with an A or a C. This was followed by a control run, during which randomly orientated gratings were presented in similar 5 stimulus sequences. C) Stimulus evoked activity of neurons responsive to the A stimulus during the control run (gray; when in the 1st, 2nd, 3rd, or 4th position) or the test run (blue). D) Stimulus evoked activity of neurons responsive to A, B, or C stimulus when that stimulus was deviant (red) or in the 4th or 5th position in the control run (gray). Magnitudes reflect standardized Caiman-estimated firing rate (eFRstd), scaled to average of control response (0-500ms) for comparison across plots. E) Bipolar electrodes inserted into V1 (200-400µm below surface) recorded LFP during the same paradigm. F,G) Morelet wavelet transformed stimulus induced power, averaged across theta (4-12Hz) and gamma (20-40Hz) frequencies for the same conditions as in C and D. One observation per mouse. Linear mixed effects model t-statistics for neurons, or paired t-tests for LFP *p<.05, **p<.01, ***p<.001.

## Results

We carried out two-photon calcium imaging with GCaMP6s in putative excitatory pyramidal neurons in layer 2/3 of awake mouse V1 (Figure 1A). Mice viewed short sets of five consecutive full-field 500ms oriented moving gratings (100% contrast, 0.08 cpd, 2cycles per second; 500-600ms ISI) separated by a 6 second inter-sequence interval. Mice first viewed 35 repeats of AAAA-B (training run, see figure 1, S1), followed by a test run, in which B was replaced by C or A every one in five sequences. After this test run, a random sequence control run was presented, lasting 70 sequences of randomly chosen orientations (0 to 157.5 degrees in 22.5 deg steps). This process was then repeated (train, test, control), swapping A and B so that the new common sequence was BBBB-A.

As previously described, we focused analyses on neurons showing statistically significant on-responses (0-500ms) to a given stimulus orientation (A, B, or C) in the deviant or the control condition (26% to stimulus A; 25% to stimulus B; 33% to stimulus C). We equated trials across conditions and, for the control condition, focused only on A, B, or C orientations appearing in the 4^th^ or 5^th^ position during the control sequence.

Responses to stimulus “A” were strongly attenuated when it was in positions 1 through 4 compared to the random orientation control (Figure 1C), indicating stimulus specific adaptation as previously reported^4,12^. Interestingly, when an A-deviant replaced the predictable B-deviant during the test run, responses to this globally deviant/locally redundant “A” were suppressed, evincing no global deviance detection (Figure 1D; left; n=94 stimulus responsive neurons from 10 mice [6 female], linear mixed-effects model t(92)=-4.06, p=7.24e-5).

However, we also did not find evidence of true deviance detection to the predictable B-deviant in the sequence (Figure 1D, middle; n=88 responsive neurons, t(86)=0.015, p=.988). While responses to B were not adapted like they were to “A”, they were not stronger than responses to B when B appeared in random stimulus sequences (during the control run).

On the other hand, when a different global deviant (“C”) was presented, which was not adapted by the preceding A stimuli (i.e., 45 degrees), or by the global context (the predictable B-deviant), a strong and significant deviance detection signal was present, relative to the control run (Figure 1D, right; n=121 responsive neurons; t(119)=2.89, p=.004).

This same pattern of effects was present in LFP recordings in V1 in a separate set of mice (n=9, 5 female; Figure 1E,F; factorial ANOVA on theta (4-12 Hz) & gamma (20-40 Hz) power^15^, 75-600ms post-stimulus onset, contextXstimulus interaction, F(3,32)=4.35, p=.011), again with significant deviance detection signal to the “C” global deviant (t(8)=2.35, p=.046) but not the “B” local-only deviant (t(8)=0.65, p=.532). Deviance detection was also absent the A stimulus was in the 5^th^ location in the LFP (globally deviant; t(8)=0.77, p=.460). Interestingly, stimulus specific adaptation was also absent to this A, in contrast to the single neuron effects (Figure 1C). Effects did not differ between sexes.

Thus, deviance detection was present only to stimuli that were neither locally redundant (the “A”) nor globally predictable (the “B”). Interestingly, during the initial training run (35 presentations of AAAAB or BBBBA), deviance detection responses were, in fact, present to the B in the first ≈7 trials, but this attenuated rapidly and was completely absent by the end of the training (Figure S1). Similar rapid contextual learning in V1 has been previously reported in spatial navigation tasks ^16,17^. Conversely, stimulus specific adaptation to A stimuli was present throughout (Figure S1).

Next we examined the distribution of responses of individual neurons to each of the deviant types (Figure 2A-F). Past reports show that deviance detection, like sensorimotor prediction errors, is not expressed ubiquitously throughout layer 2/3 neurons, but in a subset of mainly pyramidal neurons^4,5,18,19^. In line with this finding, we found that about 10.2% of all active pyramidal neurons showed deviance detection to stimulus “C” (or ≈18.3% of all neurons responsive to C; Figure 2F; one-tailed paired samples t-test with p<.025 between control and deviant, across trials), consistent with past estimates for passive visual prediction errors. These estimates were considerably lower to A (global deviant, local redundant; 3.3%) and B deviant types (global redundant, local deviant; Figure 2B,D; 3.3%)), and were visually scarce upon inspection (Figure 2A,C,E).

**Figure 2:**
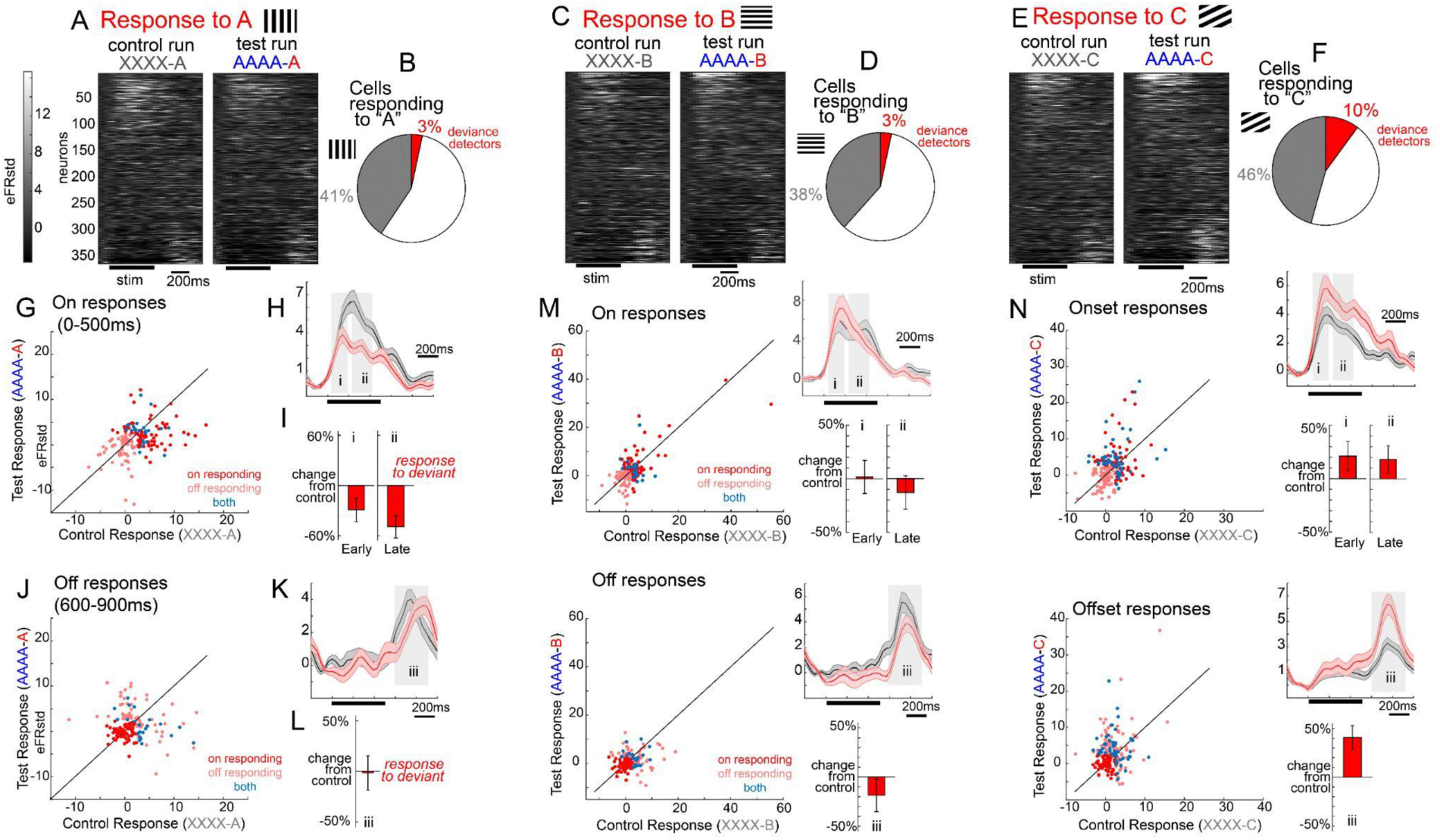
Global context impacts early and late latency onset responses as well as offset responses in V1. A) Average responses in all recorded neurons to A oriented stimuli. B) Proportion of neurons responsive to the orientation (gray) and statistically significant deviance detection (red; one-tailed t-test p<.025). C,D) same as A,B, but for the B orientation. E,F) same for C orientation. Neurons are aligned across all raster plots for cross-condition comparisons. G) Scatter plots of average on responses (0-500ms) to stimulus A when it was in the control (x-axis) vs deviant context (Y-axis). All neurons are displayed, with significantly responsive neurons colored according to the legend. H) Same as figure 1D, but indicating time windows for I) barplots of on-responsive neurons scaled to control response, for early 0-175ms (i) and late 210 to 350ms (ii). J-L) same as G-I but for off-responses. M and N) same as G-L but for stimulus B and C.

We next split neurons into on-responsive (Figure 2G-I; 0-500ms) and off-responsive neurons (Figure 2J-L; 600-900ms). Stimulus responsive neurons exhibited either only on-responses (33%), only off-responses (30%), or responses to both (37%; Figure 2A,C,E). For onset responses, about 53% of responsive neurons exhibited responses to only one orientation, while only 11.5% exhibited responses to all three. This selectivity was less pronounced for off responses, for which only 34% of PYR neurons showed selectivity for a single orientation (for a full characterization, see Figure S2). As described above, on-responses to the deviant A were attenuated, suggesting dominant stimulus specific adaptation and the absence of global deviance detection. This effect was present for both early (0-175ms) and late (210-350ms) stimulus periods (early-Cohen’s *d*=-0.21; late-*d*=-0.55). On the other hand, no such suppression was present in offset responses, nor was there significant deviance detection, to the final A (figure 2J-K; off-*d*=-0.13). To the B-deviant (i.e. the “predictable” deviant), deviance detection was absent (Figure 2M) and this pattern held true for early (*d*=0.03) and later (*d*=-0.09) response windows, compared to the control condition. On the other hand, off-responses to B were attenuated relative to the control (figure 2M, bottom; *d*=-0.33). Conversely, deviance detection to the C-deviant was present in early (*d*=0.24), late (*d*=0.27), and offset time windows (*d*=0.48). The fact that this is present early suggests that deviance detection to the C stimulus is not necessarily fed-back from higher brain regions, but is present in early spiking onset responses to the deviant C, in line with the notion of ongoing predictive modulation, occurring at – or prior to – the onset of stimuli^4^. Still, the overall pattern of offset responses suggests even more dramatic global context effects at these later latencies, as A and C offset effect sizes (Figure 2Kii) were augmented relative to the peri-onset effects (Figure 2Hi), while B responses were statistically attenuated. This suggests that later occurring feedback may additionally enhance global context processing beyond the modulation that is present at the initial stimulus evoked response, as suggested by Furutachi et al ^16,17^.

The most common paradigm for studying context processing and predictive coding in sensory areas is the standard oddball paradigm, involving randomly inserted deviant “B” stimuli in a train of redundant “A” stimuli (Figure 3A). Past work ^4,5,12,13^ shows that deviant B stimuli induce a strong response in V1 that is distinct from stimulus specific adaptation, both in magnitude (compared to a random sequence control)) and in mechanism (dependent on top-down feedback, local interneurons). However, in this standard oddball paradigm, deviant B stimuli are both local oddballs (i.e. different from the immediately preceding stimuli) and global oddballs (differing from global probabilities), making it unclear still whether deviance detection responses to B stimuli reflect local or global context^8^. In a subset of mice, we directly compared responses to B stimuli (0 or 90 degrees) when they presented in a standard oddball vs a many-standards control sequence, wherein B stimuli were equally probable as they were during the oddball run, but not contextually deviant, as multiple orientations were presented without any repetition or structure.

**Figure 3.**
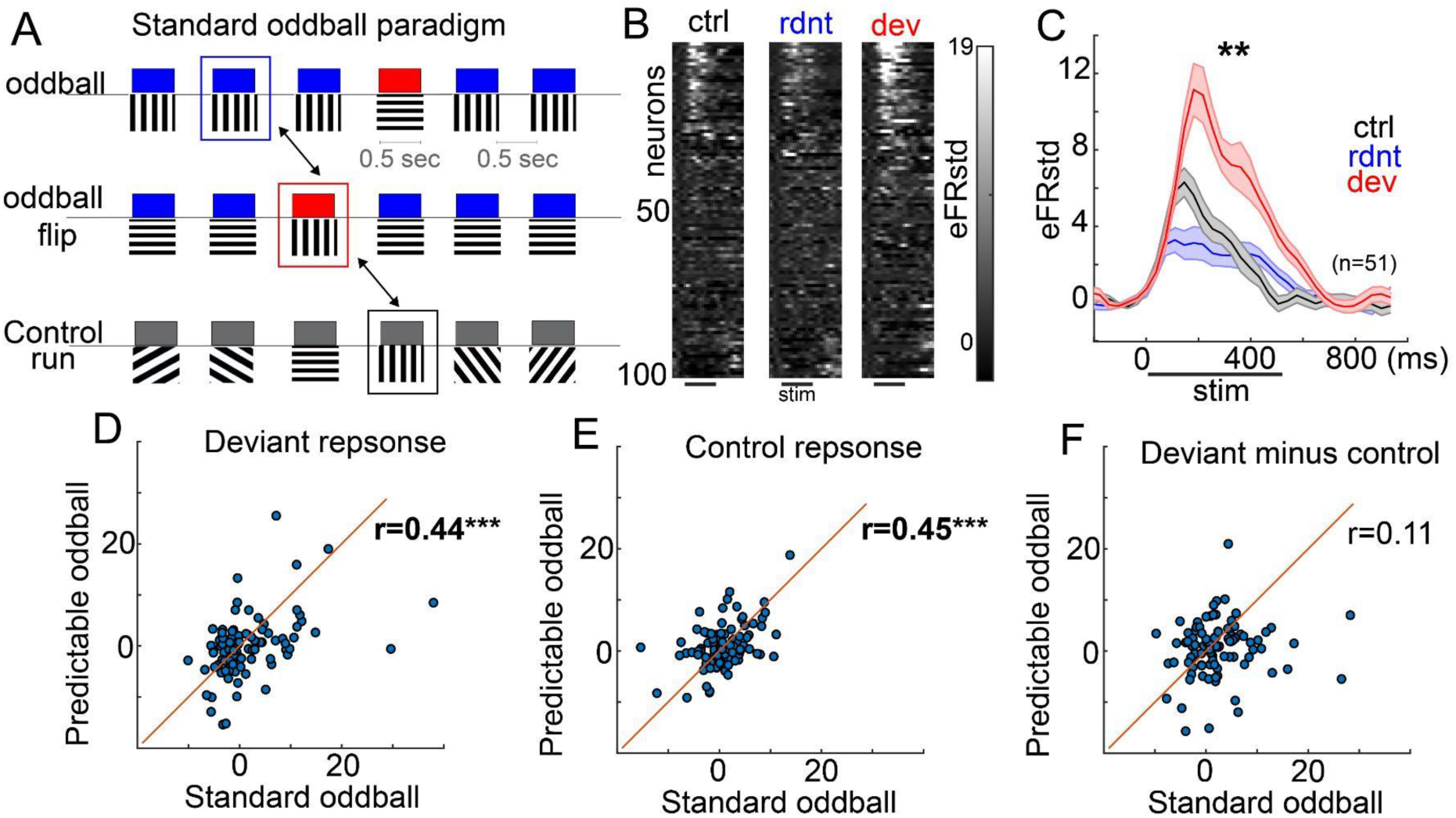
Deviance detection responses during the standard oddball reflect global context processing. A) Schematic of standard oddball paradigm, using the same features, duration, and ISI as the global/local, but continuous presentation and no predictability of oddball timing. Comparisons are made to a many-standards control sequence (bottom) where 1 of 8 oriented stimuli are presented at random, such that each orientation (stimulus B) is equally probable as in the oddball run. B) Average responses of recorded V1 layer 2/3 neurons from 3 mice to the same orientations when it was neutral, redundant, or deviant. C) Stimulus evoked activity of neurons responsive to one of the two oriented stimuli (n=51). D) Comparing responses in each neuron to the oddball (deviant) orientations during the standard oddball (x-axis) vis the predictable oddball (i.e., the B when AAAAB was trained). E) Same as D but for the control run. F) Same as F, but for the computed deviance detection score of deviant minus control. Pearson correlations, ***p<.001.

As expected, B responses in the standard oddball were dramatically larger than they were to the control, suggesting deviance detection that was distinct from simple stimulus specific adaptation (Figure 3B,C; responsive neurons, t(49)=2.99, p=.003). When examining the same neurons responding to “B” during the standard oddball vs the predictable (global/local) oddball, response magnitudes were correlated between the two deviant (r=0.44; p<.0001) and two control runs (r=0.45, p<.0001) as expected. However, these standard oddball deviance detection responses (deviant minus control) were considerably larger than the responses in the same neurons to the deviant-B stimulus when it was *globally* predictable (i.e., during the AAAA-B sequences; all neurons t(98)=2.14, p=0.033) and, curiously, were not correlated (r=0.11; p=0.276) suggesting that deviance detection in the standard oddball is distinct from the local context modulation present in the global/local paradigm.

In sum, the standard oddball paradigm, used for decades to study basic sensory-cognitive function in health and neuropsychiatric disease^20,21^, evokes deviance detection responses that reflect at least some degree of higher-order context processing. In the auditory system, past work has shown that randomly timed oddballs elicit larger responses than periodic (temporally predictable) oddballs^22^ – a point that is in line with our contrast between the standard oddball and the local AAAAB oddball. Critically, isolating this deviance detection requires the use of appropriate contrasts in the analyses (i.e., deviant vs control, rather than deviant vs redundant).

Finally, past work has shown that basic deviance detection in early sensory areas during the simple oddball paradigm depends on top-down feedback to V1^4,5^ or A1^7^. Here we tested whether such feedback is also necessary for global context modulation seen during the global/local paradigm. We focused on anterior cingulate area (ACa), which has previously been shown to densely innervate V1 and support context processing in a variety of paradigms^4,23–25^. We targeted this feedback circuit using a viral strategy (Figure 4A), expressing inhibitory DREADDs receptors (hM4Di) in excitatory neurons of ACa, followed by electrode implantation and delivery of DCZ (a DREADD ligand) or saline on separate days (Figure 4B,C). LFP recordings in a subset of mice confirmed DREADD-based suppression of ACa activity (high-gamma, 80 to 200-Hz power; LME with order (day 1 vs 5) as random effects factor, t(14)=-3.24, p=.006) but not V1 (t(14)=-1.27, p=.223; Figure 4D,E). This suppression of ACa abolished contextual modulation in V1 during the global local oddball paradigm (LME with order as random effects; 3-way interaction of drug by context by deviant-type, t(87)=-2.19, p=.031). On the day with saline injection, mice exhibited clear deviance detection to the global C deviant (Figure 4F, t(11)=-3.26, p=.008) but not the predictable B deviant (t(11)=1.299, p=0.22). On the day with DCZ, mice did not exhibit deviance detection in any condition (figure 4G). Further, we quantified the degree to which mouse V1 responses were augmented to global deviants (stimulus C) relative to local-only deviants (stimulus B). After saline, V1 responses showed robust augmentation to C relative to B, and this was strongly attenuated after DCZ (Figure 1H). Likewise, ACa suppression also appeared to modestly augment responses to the control as well (Figure 1I). This bidirectional modulation is consistent with past findings during the oddball paradigm^4,5^, suggesting not a simple top-down inhibition or augmentation of V1 during visual processing, but a context-dependent modulation of response gain.

**Figure 4.**
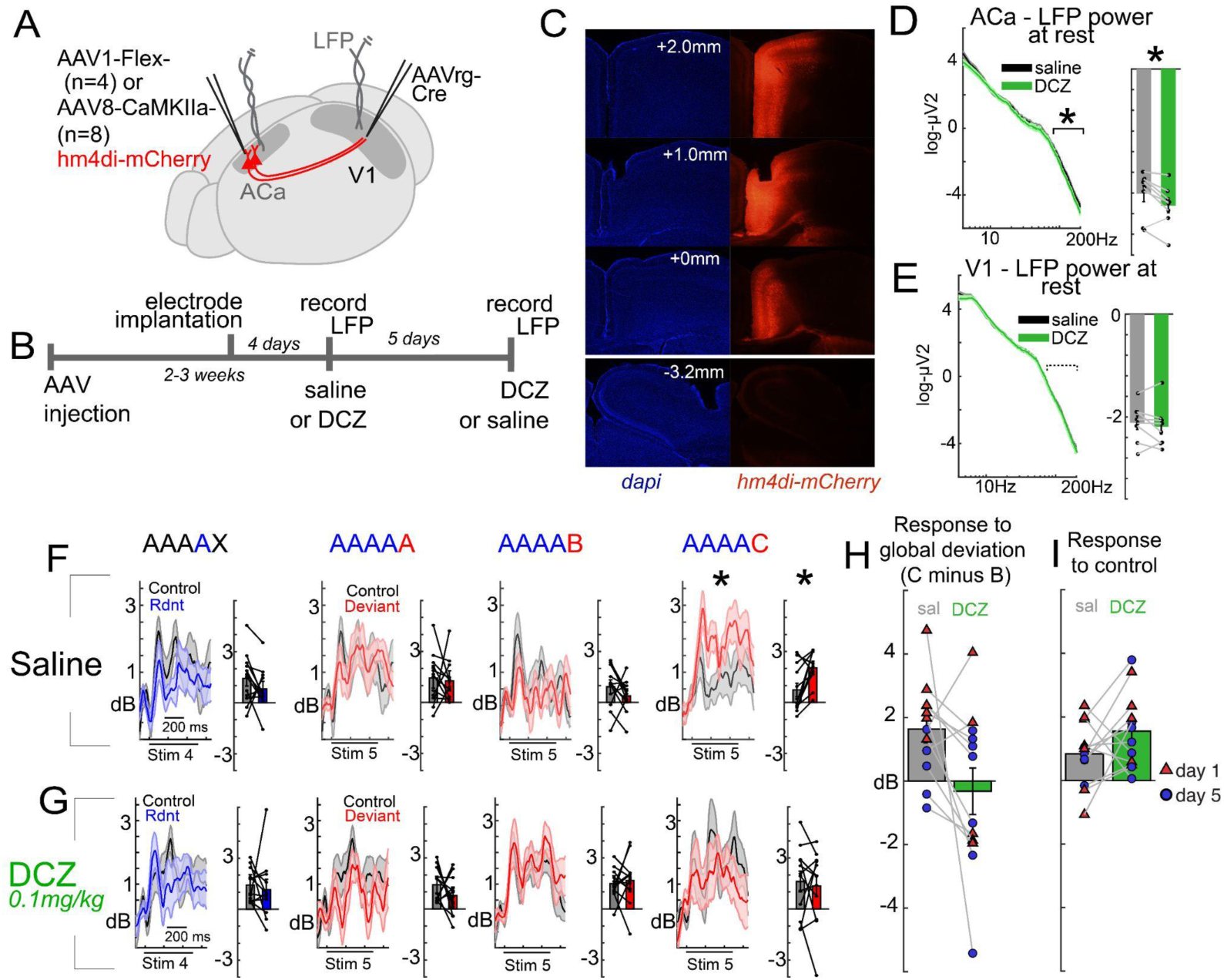
Global context processing depends on top-down feedback to V1. A) AAVs targeting ACa to V1 projections with inhibitory DREADDS (hm4Di) were injected into ACa and bipolar electrodes were inserted into ACa and V1 in 12 mice. B) Two weeks later, local field potentials were recorded during the global/local oddball paradigm after mice were given 0.1mg/kg DCZ (to inactivate ACa to V1 projections) or an equivalent volume of saline I.P. 5 days later, mice were given the other treatment (saline or DCZ) and LFPs were recorded again. C) Electrode location and hm4Di expression were confirmed post-hoc. D) DREADDs effectively decreased high gamma LFP power in ACa (80 to 200 Hz; n=8 mice, subset with confirmed ACa electrodes) but E) not V1. F) After saline injections, mice exhibited global deviance detection to C but not B, replicating figure 1F (theta/gamma power, 75 to 400ms post-stimulus onset). G) When ACa was suppressed with DCZ, contextual modulation during the paradigm was absent. This effect was exhibited in H) reduced global vs local oddball responses after DCZ (C in AAAAC minus B in AAAAB) and I) increased responses during the control run (to both C and B combined).

## Discussion

These results demonstrate that global context *can* modulate early visual cortical activity, yet provide mixed support for predictive processing theories. Prediction error-like responses were absent in V1 to globally predictable B stimuli, but they were present to both the globally and the locally deviant C, in support of predictive coding theories. The higher-order context – that a B was predictable at the end of each sequence – was learned rapidly, within the first exposure session, and modulated early visual cortical responses at the neuron level. Context dependent response modulation to C vs B in these sequences was dependent on higher cortical areas that feedback to V1, namely, ACa. This point is also consistent with predictive coding theories.

However, although deviance detection was absent to predictable-B stimuli, responses to predictable-B were also *not suppressed* relative to the contextually neutral-B. Likewise, during chemogenetic suppression of ACa, deviance detection responses to the predictable B did not reemerge. These findings argue against a top-down predictive *suppression* in V1, conflicting with classic predictive coding theories. Instead, we propose that top-down feedback in predictable contexts may ultimately play a gain modulation role, amplifying responses to deviants, rather than suppressing responses to redundants – closer to a top-down disinhibition model^3,4,26^.

And, perhaps most starkly in contrast to predictive coding theories, standard deviance detection was absent to the globally deviant-A in V1 – at least in the firing outputs of layer 2/3 neurons. We propose an update to predictive coding ideas wherein bottom-up adaptation dominates V1 responses in contexts where there is strong redundancy, such as an “A” deviant which comes after a string of repeated “A” stimuli, and thus circumvents the calculation of prediction errors at this early cortical region. Past work has shown that signatures of stimulus specific adaptation are present both at earlier latencies and further upstream than signatures of deviance detection^13,14^. It is possible that if certain stimulus features are already adapted sufficiently prior to reaching deviance detection computing circuitry in layer 2/3, deviance detection may be dampened or altogether eliminated in V1. That said, another possibility is that mice do not perform true counting^27^. That is, the internal model learned in this paradigm could be that of a B coming after some approximate but unquantified string of A stimuli. Therefore, an A in the 5th spot does not trigger a prediction error, because the mouse cortex does not differentiate the fifth A from the 4th A. Future work could test this by inserting a B stimulus into the 3rd or 4th position. A deviance detection response to the B in the 4th or 3rd position, but not the 5th, would clarify the nature of the internal predictions.

Interestingly, our experimental results replicate those of a recent study by Westerberg et al, though our overall conclusion differs, with our interpretation informed by our extended experimental design. We find that higher level context modulation and complex deviance detection *is* present in mouse V1. The Westerberg study reported augmented responses to the local oddball (i.e., the globally predictable B in the learned AAAB sequence) when compared against the third “B” in a BBBB control condition, and ascribed that to a true deviance response. Ostensibly, our data are also consistent with this pattern (see figure 1C vs D), but we argue that a more appropriate contrast is a “B” stimulus present under control conditions where orientations are mostly unpredictable – and which are not preceded by a B stimulus (see Methods: Quantification and Statistical Analysis). This directly avoids the stimulus specific adaptation that would be present under a BBBBA condition.

It is possible that longer-term adaptation effects in V1, emerging across days, could have altered responses in the Westerberg study, as previously shown^28^. In Westerberg et al., mice viewed the sequences for three to five days, over two thousand times, prior to testing. In our study, mice saw the sequence only 35 times during a single habituation sequence just prior to testing, and we found that deviance detection to the globally predictable “B” was eliminated within *minutes* of the initial habituation sequence. This is consistent with local field potential studies, where, within a single session, signals in mouse V1 demonstrated complex sequence learning (e.g. ABCD) and deviance detection (e.g. ABDC)^11^.

While mounting evidence across studies calls into question a simple and complete application of classic predictive coding, one possibility is that predictive processing applies well to only a subset of neurons or a sub-circuit within the cortex (e.g. only ≈12% of our neurons showed deviance detection to the C in figure 2), while other embedded non-prediction-modulated circuitry supports a) the maintenance of feed-forward processing streams^29^ (regardless of predictability) and/or b) top-down attentional modulation (which, in contrast with classic predictive coding, can serve to enhance responses to anticipated stimuli in some cases^30^).

Notably, our single neuron calcium imaging recordings suggest that deviance detection responses (to the global and local oddball C) are present in V1 at the earliest latencies, consistent with our past reports in simple oddball paradigms^13^. A recent theoretical paper ^31^ argues that such early latency deviance detection responses (even to complex sequences, as reported in^11^) are evidence that they reflect local adaptation that is simply fed-forward from lower levels, rather than integrative prediction errors. However, another possibility explaining this early latency effect is that top-down modulation in predictable contexts is ongoing/tonic and present in anticipation of visual stimuli, i.e. prior to or timed to the *onset* of anticipated stimuli^4^. In such a manner, it modulates V1 not in a reactive manner on a trial by trial basis, but in an ongoing anticipatory manner, altering excitability with respect to contextually typical features (see^3^).

Consistent with this point, feedback inputs to V1 from mouse ACa do not themselves exhibit deviance detection during standard oddball paradigms, as evidenced by calcium imaging of ACa axonal segments in V1^4,5^, despite the fact that such feedback inputs are nevertheless necessary for deviance detection within V1. This suggests that ACa supports deviance detection not by relaying the deviance detection signal back to V1, but by modulating V1 in an ongoing, tonic fashion to alter context-dependent excitability. In our experiments, ACa’s role in modulating V1 was not simply inhibitory or excitatory, but was modulatory – in some cases, ACa suppression enhanced V1 activity (Figure 4I) while in others, it suppressed activity (Figure 1G, H).

Future work should determine how these modulatory signals interact with local inhibitory microcircuits in V1 and whether disruptions of this modulatory circuitry contribute to altered predictive processing in neuropsychiatric disease^32^. The results presented in Figure 3 carry implications for interpreting a volume of clinical studies on sensory integration in psychiatry^33^. The basic oddball paradigm is ubiquitous in clinical neuroscience research, as several large scale patient population studies employ simple oddball sequences for extracting EEG biomarkers of deviance detection^34,35^. Our findings emphasize that the deviance detection signals evoked during these simple paradigms include aspects of higher order cognition – a point consistent with the fact that simple oddball paradigms correlate with multiple higher level symptoms and cognitive deficits in schizophrenia^33,36^.

## Acknowledgments

This work was funded by the National Eye Institute (R01EY033950, Hamm), National Institute of Mental Health (F32MH125445, Ross; R01MH128176, Hamm), and the Brain and Behavior Research Foundation (YI30149; Hamm).

## Author Contributions

J.P.H. D.S.P., and F.I. designed research; F.I., G.B., J.R., L.R, M.H, and J.P.H. performed research. J.P.H. analysed data. All authors wrote and edited the paper.

## Competing interests

The authors declare no competing interest.

## Methods

### EXPERIMENTAL MODEL AND STUDY PARTICIPANT DETAILS

Experiments were carried out under the guidance and supervision of the Georgia State University (GSU) Division of Animal Resources and the Nathan S. Kline Institute (NKI) Division of Animal Resources. All procedures were approved via Institutional Animal Care and Use Committees (IACUCs) at GSU and NKI. Adult C57BL/6 mice (n=33, from Jackson Laboratories) were used. Mice were weaned at P25 and group housed (max 5 mice per cage) separated by sex. They were maintained at a 12/12 light/dark cycle (light: 6am-6pm) with food and water available ad libitum. Animal ages varied between P73(min) and P132 (max) and averaged at P99 days. Experimental mice were chosen randomly based on availability and age; both female and male mice were included in similar proportions (16F, 17M). For calcium imaging, transgenic lines were made using mice expressing cre-dependent GCaMP6s (tm162(tetO-GCaMP6s, CAG-tTA2)) crossed with tm1.1-(VGluT) lines. For LFP recordings, we used wildtype mice of C57Bl/6 origin.

#### Surgical procedures

For experiments involving calcium imaging, head-plate fixation and craniotomy surgeries were carried out together as previously described ^4,5,12,37^. Briefly, a hole with a 3 mm diameter was drilled in the mouse skull in left V1 (coordinates from bregma: X = 2 mm, Y = −2.92 mm), followed by the removal of the skull and exposure of brain surface; dura matter was conserved. A cover glass was placed and sealed at the hole location. Then a titanium head-plate was attached to the mouse head to allow for head-fixation during imaging.

For ACa suppression experiments, we either a) injected AAV1-hSyn-flex-hM4Di-mCherry (Addgene #44362, 2.4×1013) in ACa and AAVrg-hSyn-Cre (Addge #105553, 2.5×1013) in V1 (n=4) or b) 300nl of AAV8-CaMKIIa-hM4Di-mCherry (Addgene #50477 2.1×1013) in ACa (n=8) stereotaxically. Following coordination was used. ACa: Two sites (200nl each), coordinates from bregma: x = 0.3 mm, y = 0.6 and 1.2 mm, z = 0.9 mm), V1: 300nl, coordinates from bregma: x = 2.2 mm, y = −2.9mm, z = 0.5mm). After two weeks, bipolar electrodes (2 Channel Electrode, P1 Technologies, #MS303/2-A/SPC) were implanted.

For local field potential experiments, head-plate attachment was carried out prior to electrode implantation. A bipolar electrode (with contacts spaced approximately 200 µm apart) was inserted below the dura in stereotaxically defined V1 (coordinates from lambda, X =2.3mm, Y = 0.7mm) approximately 0.5mm between the dura and white matter. For DREADD experiments, to confirm the impact of DREADD suppression, mice had a second electrode inserted into ACa (Coordinates from bregma, x=0.3mm, y=1.0mm, z=0.9 mm). In four of these mice, ACa targeting was unsuccessful, totalling 8 for statistical comparisons (figure 4D). Prior to recordings, mice underwent multiple training sessions to acclimate them to head-fixation, as previously described ^4^. Mice only viewed the oddball paradigms on the day of recording. On the day of recording, mice first viewed the regular oddball paradigm, followed by the global/local paradigm

### METHOD DETAILS

#### Visual Stimulation

Visual stimulation was presented on a 27 inch flat LCD screen (Dell, P27P5) at a 45° angle from the animal axis, approximately 15 cm from the eye, using Psychophysics Toolbox on MATLAB (Mathworks). Stimuli were full-field, black and white, sinusoidal moving gratings were presented at 100% contrast, 0.08 cycles per degree, two cycles per second, at 8 possible orientations (30°, 45°, 60°, 90°, 120°, 135°, 150°, and 180°).

For the global/local paradigm, short sequences of five stimuli of 500ms in duration, with an inter-stimulus interval of 500-550ms of gray screen, were presented, followed by 6000ms of gray screen (inter-sequence interval). Initially, mice viewed a “training set”, which included 70 sequences of AAAAB where A was 0° and B was 90°. After this, mice rested for 2-3 minutes, and then viewed a “test set”, wherein AAAAB sequences were primarily presented, but AAAAA (global deviant, local redundant) and AAAAC (global deviant, local deviant) catch trials were presented one in 6 presentations (C was always 45 °). Then, after another brief rest, a control run was presented, wherein random orientations (e.g. CDEAB) were presented in the same 5 stimulus sequences (500ms duration, 500-550ms ISI, 6000ms inter-sequence interval). A subset of mice then rested 5 minutes and this set of three paradigms was repeated, where A and B were switched.

For the standard oddball, the same stimulus duration and ISIs were used, but stimuli were presented in continuous trains of 250 trials (≈4 minutes). This consisted of a “many-standards control” (equally rare, randomly presented stimuli at all 8 possible orientations) and an oddball sequence. The oddball sequence consists of a repetitive sequence of one stimulus (“redundant”, either 0° or 90° degree angles, presented 87.5% of the time), randomly interrupted by a stimulus of a different orientation (“deviant”, 120°, 135°, or 180° degree angles, presented 12.5% of the time). In the latter half of the sequence, the redundant stimulus is “flipped” to become the deviant, and vice versa (“oddball flip”).

#### Two-photon calcium imaging

Two-photon microscopy (28 Hz framerate; Bruker Investigator laser scanning two-photon microscope; Bruker Corporation, Billerica, MA, USA) excited by a laser (Chameleon Ultra II, Coherent Inc, Santa Clara, CA, USA) at 920 nm wavelength was used to image the fluorescent calcium sensor GCaMP6s expressed in excitatory neurons in the mouse visual cortex. The laser beam was modulated with a Pockels cell (Conoptics 350-105, with 302 RM driver) and scanned with galvometers through a water immersion objective (16X/0.80W, Nikon, Tokyo, JP). The objective lens was positioned over the animal’s head while a small volume of Aquasonic ultrasound gel (Parker Laboratories Inc) was placed at the site of the cranial window to bridge the objective with the imaging area in the sessions. The animals were awake, head-fixed under the microscope by their headplate, while sitting on top of a treadmill (low-friction mouse belt-treadmill with rotary encoder, from LABmaker, Berlin, Germany) free to move forward and backwards. All recordings were carried out in a dark room with the researcher present to monitor mouse wakefulness and check for signs of discomfort. Each run had a duration between 4-13 minutes. Scanning and imaging were done through Prairie View (Prairie Technologies) software (8 KHz resonant galvanometer, recorded at ~56 frames per second, downsampled to 28 frames per second by averaging two-frames, for 256×256 pixels, 3.136 µm pixel size, 802.9 x 802.9 µm field of view). A time-series was recorded using Prairie View software as the mice observed visual stimuli. The visual stimulus was transmitted to the monitor through an HDMI cable, converted to a voltage trace, and connected to the computer through the Voltage Recording tool on Prairie View. Time-series and stimulus voltage traces were synchronized at the onset of recording for proper alignment of neuronal activity and stimulus presentation. Images were taken 100 µm to 350 µm from the brain surface, approximately L2/3 of the mouse cortex. Locomotion was recorded during two-photon calcium imaging via a rotary encoder integrated into the treadmill.

#### Local field potentials recordings

For the LFP experiments, the mouse was fixed to the recording apparatus through the head-plate and free to locomote on the treadmill. Insulated cables were connected to the implanted electrodes and plugged into a differential amplifier (Warner instruments, DP-304A, high-pass: 0 Hz, low-pass: 500 Hz, gain: 1K, Holliston, MA, USA). Amplified signals were passed through a 60 Hz noise cancellation machine (Digitimer, D400, Mains Noise Eliminator, Letchworth Garden City, UK), which, instead of filtering, creates an adaptive subtraction of repeating signals which avoids phase delays or other forms of waveform distortion. Prior to insertion, probes were submerged in DiI dye for post-hoc anatomical validation. Electrophysiological activity was recorded using the Prairie View software. Visual stimulus timings were recorded as voltage traces at the same time as the LFPs signals for proper alignment. Locomotion was recorded during electrophysiology via a rotary encoder integrated into the treadmill.

#### Chemogenetic suppression of ACa

On the day of recording, mice were injected with DCZ (Sigma-Aldrich, SML3651, final concentration 0.1mg/kg)^38^ or saline I.P and fixed to the treadmill. Ten minutes later, mice viewed regular oddball paradigms followed by global-local oddball and control runs as described above (one set of each) and then returned to their home cage. Four days later, they were injected with the opposite drug (i.e. saline if they received DCZ on day 1, or vice versa) and recordings were repeated.

#### Histological verification

Mice were deeply anesthetized via urethane and 3% isoflurane and then perfused transcardially with PBS followed by 4% paraformaldehyde (PFA)/PBS. Heads with electrodes and headplates attached were post-fixed in 4% PFA/PBS for two overnights. After isolation, brains were cryoprotected with 30% sucrose/PBS for one to two overnights and then embedded in Tissue-Tek OCT compound (Sakura) and cryosectioned (50 µm). Sections were stained with DAPI (ThermoFisher, D1306, 1µg/ml) and mounted using antifade medium (VectorLabs, H-1700-2).

### QUANTIFICATION AND STATISTICAL ANALYSIS

#### 2-photon image processing and analysis

Calcium imaging data sets were scored similarly to previous reports^37^. Raw images were processed to correct motion artifacts using the “Moco” plugin for ImageJ (NIH)^39^. Regions of interest (ROIs) were selected semi-automatically from time-averaged images using an in-house written MATLAB (MathWorks) routine and fluorescence traces were subtracted from the pixels just outside these ROIs (i.e., halo subtraction). Because of the relatively rapid stimulus presentation used in this study (≈1Hz) relative to the decay time of GCaMP6s, neural activity was analyzed by deconvolving putative spikes from fluorescent traces using a constrained non-negative matrix factorization algorithm (constrained-foopsi^40^, available in the Oasis or Calman toolboxes^41^) and smoothing the result with a 140ms gaussian window (5 samples) to estimate firing rate (eFR). For analyses, eFR was standardized (eFRstd) using the standard deviation of the bottom 50% of positive values across the entire recording. Standardizing did not impact the statistical conclusions or the pattern of the effects, but helped normalize values for clearer visualization across neurons.

For the global-local paradigm, eFRstd was averaged across trials for each stimulus type (control-A,B,C; deviant-A,B,C; redundant-A). As each mouse saw just 6 of the type A and C deviants, we limited analyses to 6 trials of each of these conditions. For the control run, we only analyzed A, B, or C orientation trials i) if they fell in the 4^th^ or 5^th^ spot in the sequence, and ii) if they were not an immediate repeat (i.e., a C stimulus in the 3^rd^ and 4^th^ spot). For A and B stimuli during the testing run, we analyzed trials in the sequence immediately preceding the catch sequences. That is, the 4^th^ A or 5^th^ B response in AAAA-B that comes right before an AAAA-C. For the standard oddball paradigm, we also limited to 6 trials in order to facilitate optimal interparadigm comparisons. We focused on the first 6 deviants of each orientation and the final 6 instances of a control orientation during the control run (as the control sequence was always played first, this helped keep the trials being compared closer in time). Further, controls were excluded if they were repeats with the previous trial, as described above. Further, trials with locomotion were excluded (<15% of trials for any mouse).

As previously described, we focused analysis on cells with significant stimulus-evoked responses to a given orientation i) average responses >1.97 standard deviation (p<.05, two-tailed) or ii) average responses >2.6 (p<.01) above its pre-stimulus baseline on both the control and the deviant condition of a given orientation. Importantly, these choices of cell and trial inclusion had no tangible effects on the conclusions, but were chosen to allow for the most accurate effect size estimates and consistency with past studies.

#### Local field potential signal processing and analysis

Trials with excessive signal (>≈5 std devs) in V1 were manually excluded (between 0 and 20). All analyses were limited to the same trial choices as described above. Ongoing data were converted to the time-frequency domain with a modified morelet wavelet approach with 79 evenly spaced wavelets from 4 to 100Hz, linearly increasing in length from 1 to 20 cycles per wavelet, applied every 5ms from 300ms pre- to 700ms post stimulus onset (200ms post-stimulus offset) as previously described^2^. Stimulus-induced power spectra were computed for all conditions for each mouse and baseline-corrected by subtracting the average for each frequency in the 100ms prior to stimulus onset. Based on past work ^4,13^, we compared control vs deviant responses in theta (4-12Hz) and gamma power (20-40Hz) and the 75 to 600ms time-period. For analysing whether DREADDs impacted ACa, we analysed baseline power. We segmented LFP recordings during the control and oddball runs, focusing on 1 second periods with no locomotion or visual stimulation. We equated the number of segments across all mice (n=245). We carried our fast fourier transforms, taking the natural log of the power (squared amplitude) for each segment, and then the median value for each frequency (4 to 200 Hz) across segments, for each mouse, condition (Sal, DCZ), and region (V1 vs ACa). Analysis focused on high gamma power, 80 to 200-Hz. Histological verifications indicated that all mice except one had V1 successfully targeted. Additionally, 8 mice had ACa success

#### Statistics

For calcium imaging analyses, we focused on cell-wise analyses and carried out a linear mixed-effects model to account for the fact that many neurons came from the same mice (mouse as random-effects). In these models, significant effects of deviant vs control were the primary focus, and were reported as t-values. For LFP experiments in figure 1, only one set of values came from each mouse, so we carried out factorial repeated-measures ANOVAs on induced power at the mouse level with stimulus type (CONTROL, DEVIANT) and deviant type (redundant, (A) local-redundant, global-deviant, (B) local-deviant, global-redundant, (C) local deviant, global deviant) as within-subjects factors. Significant interactions were tested further with paired-samples t-tests. For DREADDs experiments (figure 4), we focused on deviant types B vs C, as this is where our main effects were in figures 1 and 2. As some mice received saline first while some received DCZ first, we included an ORDER variable as a random effects factor (Day 1 vs Day 5). We carried out linear mixed effects models on induced power at the mouse level with stimulus type (CONTROL, DEVIANT) and deviant type (redundant, (A) local-redundant, global-deviant, (B) local-deviant, global-redundant, (C) local deviant, global deviant) as within-subjects factors and drug condition (SALINE, DCZ) as a between subjects factors. We tested specifically for a 3 way interaction of stimulus type, deviant type by drug condition, controlling for order effects. For analysing baseline power, we used a linear mixed effects model analysing drug condition (SALINE, DCZ) while controlling for order as a random effects factor.

## Notes

### Competing Interest Statement

The authors have declared no competing interest.

