## Supplemental Figures for "Global context rapidly shapes sensory responses in V1"

Supplemental Materials

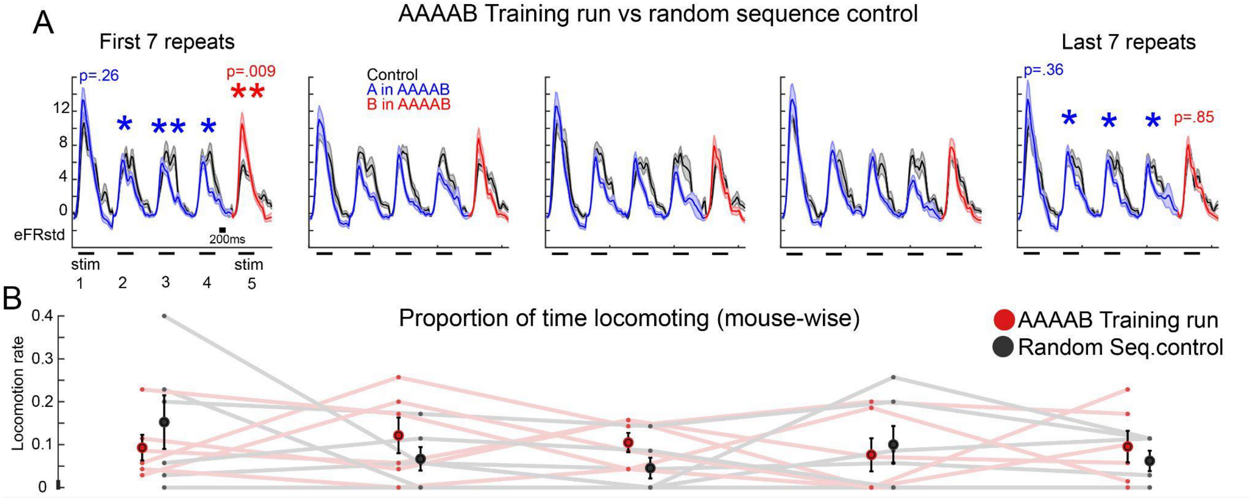


**Figure S1. Mouse V1 rapidly learns AAAAB sequence and suppresses deviance detection.** A) Trials during training sequences (AAAAB and BBBBA) and the control sequences (e.g. CDAEB) were split into pentiles of 7 trials each. Neurons active at both the beginning and end bin at $p<.05$ or active during either at $p<.01$ were included for analysis. Data come from 152 responsive neurons from 5 mice, four runs each (AAAAB training, BBBBA training, and two control runs). Trials with locomotion were not excluded in order to ensure sufficient data for all mice for all bins. Linear mixed effects models (df=150) were ran at the neuron level with mouse as a random effect. Were carried out for stimuli 1 through 5 in the first bin and the last bin. *$p<.05$; **$p<.01$. B) Locomotion did not vary as a factor of bin for control or oddball runs (bin main effect- $F(1,4)=.071$, $p=.589$; bin by context interaction– $F(1,4)=1.36$, $p=.263$).

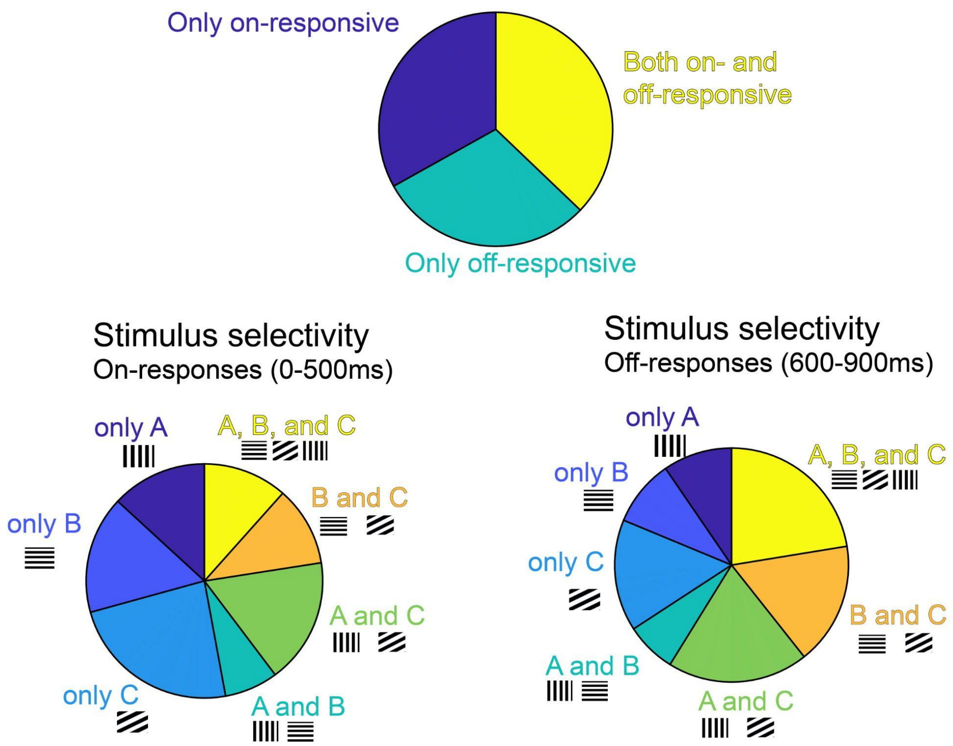


**Figure S2. Selectivity of neurons in analysis**. Selectivity determined as neurons with average responses to the stimuli and/or time windows greater than baseline at p<.01 (z-test). Top: Onset- vs Offset- vs Both selectivity was evenly split among responsive neurons. Bottom: On responses showed more pronounced feature selectivity than off responses.
